# Preference Shifts During Multi-Attribute Value-Based Decisions

**DOI:** 10.1101/2023.05.10.540218

**Authors:** Ayuno Nakahashi, Paul Cisek

**Affiliations:** Department of Neuroscience University of Montréal Montréal, Québec, CANADA

**Author notes:** Corresponding author address: Paul Cisek Department of Neuroscience, University of Montréal C.P. 6128 Succursale Centre-ville Montréal, Québec, CANADA.

## Abstract

When choosing between options with multiple attributes, do we decide by integrating all of the attributes into a unified measure for comparison, or does the comparison occur at the level of each attribute, involving independent competitive processes that can dynamically influence each other? What happens when independent sensory features all carry information about the same decision factor, such as reward value? To investigate these questions, we asked human participants to perform a two-alternative forced choice task in which the reward value of a target was indicated by two independent visual attributes – its brightness (“bottom-up” feature) and its orientation (“top-down” feature). If decisions always occur after integration of both features, there should be no difference in the reaction time (RT) distribution regardless of the attribute combinations that drove the choice. Counter to that prediction, almost two-thirds of the participants exhibited RT differences that depended on the attribute combinations of given targets. The RT was shortest when both attributes were congruent or when the choice was based on the bottom-up feature, and longer when the attributes were in conflict (favoring opposite options), especially when choosing the option favored by the top-down feature. We also observed mid-reach changes-of-mind in a subset of conflict trials, mostly changing from the bottom-up to the top-down-favored target. These data suggest that multi-attribute value-based decisions are better explained by a distributed competition among different features than by a competition based on a single, integrated estimate of choice value.

**New & Noteworthy:** This study showed that during value-based decisions, humans do not always take all information about reward value into account to make their choice, but instead can “jump the gun” using partial information. In particular, when different sources of information were in conflict, early decisions were mostly based on fast bottom-up information, and sometimes followed by corrective changes-of-mind based on slower top-down information. Our results suggest that parallel decision processes occur among different information sources, as opposed to between a single integrated “common currency”.

## INTRODUCTION

Many of our decisions are based on multiple sources of information about a wide variety of factors. For example, when choosing a house to buy, we consider its cost and size, the property and environment, as well as its location relative to workplaces, good schools, public transportation, etc. Such multi-attribute decisions also occur in the wild, when for instance, animals need to weigh the value of seeking food against the effort required to obtain it and the potential exposure to danger. One plausible model suggests that the multiple factors pertinent to each option are integrated together into a central representation of subjective value that reflects them all, and each factor is weighted by the degree to which it matters for the deciding agent (Cai & Padoa-Schioppa, 2019; Kable & Glimcher, 2009; Levy & Glimcher, 2012; Padoa-Schioppa, 2011; Shizgal, 1997). This permits comparison between options that may differ in every way, such as when choosing between a small apartment that is close to work versus a large house that will require a longer daily commute, and permits a rational decision process that optimizes some general measure of global utility. At a mechanistic level, such “integrated competition” models predict that only a single comparison is made in the brain – a comparison between the integrated values of the options under consideration.

Alternatively, “distributed competition” models of multi-attribute decision-making suggest that the individual factors can themselves compete against each other prior to integration, and even that the weights of the factors can vary over time due to fluctuations in covert or overt attention (Busemeyer & Townsend, 1993; Diederich, 2003; Hunt et al., 2014; Krajbich & Rangel, 2011; Roe et al., 2001; Trueblood et al., 2014; Yang & Krajbich, 2022). For example, Hunt and colleagues suggested that during multi-attribute decision-making tasks, human behavior is best explained by a hierarchical scheme in which the higher-level competition between choices is biased by information from multiple lower-level competitions that each compare the options in terms of a given attribute, and that the weighting of each attribute is itself subject to a competition based on its relevance and discriminatory efficacy (Hunt et al., 2014). Such models predict that even when the total value should favor choice A over B, subjects may sometimes choose B if one of its attributes is particularly outstanding. Another prediction of such models is that when decisions are made quickly, commitment can be reached even before all of the factors have been considered (Diederich, 2003), explaining preference reversals as a function of deliberation time.

Here, we ask whether the competition can be distributed even among different sources of sensory information about a single factor, such as reward value. In particular, we ask how different visual features, both of which carry information about value, are integrated into the decision process. Importantly, certain kinds of “bottom-up” visual signals are processed very quickly, including information about spatial location of stimuli and simple features like brightness. This is believed to implicate the dorsal visual stream from occipital cortex to the posterior parietal lobe (Galletti & Fattori, 2018; Goodale & Milner, 1992; Mishkin & Ungerleider, 1982; Treisman & Gelade, 1980), throughout which sensory responses have been reported in as little as 50ms (Ledberg et al., 2007; Schmolesky et al., 1998). Other kinds of visual information require more sophisticated and slower “top-down” processing, including categorization of arbitrary shapes according to task-dependent rules, which involves the ventral visual stream and prefrontal cortical regions (DiCarlo et al., 2012; Freedman et al., 2002; Goodale & Milner, 1992; Kravitz et al., 2013).

In this study, we presented human participants with a two-alternative forced choice task in which the reward value associated with each choice was determined by two different visual features – a bottom-up brightness cue and a top-down orientation rule – and asked whether the timing of their responses revealed how these features interact. Of particular interest were situations in which the two features were in conflict, such that the total reward value of each choice was identical, so it did not matter which was chosen but a choice still had to be made.

In those trials, was the choice random or did it depend on the relative timing with which each feature was processed with respect to the time of commitment? Importantly, these trials were interleaved among a variety of other trials in which one choice was clearly better than the other. Consequently, because value could only be evaluated after both features were processed, participants were motivated to take the time to integrate both kinds of information prior to committing to either choice, especially since there was no time pressure to respond quickly. Therefore, “integrated competition” models would predict that participants should always take the time to evaluate the options fully, so decision timing should be similar across all choices, whether or not they are easy (both features favor the same target) or ambiguous (different features favor different targets). By contrast, “distributed competition” models would predict preference changes as a function of time, with initial choices more biased by bottom-up stimulus features and later ones increasingly based on top-down information.

The design of our task also allowed us to examine whether the effect of reward size on reaction times and choice accuracy follow predictions from the Weber-Fechner law, which states that the “just noticeable difference” (JND) between stimuli increases with stimulus amplitude (Fechner, 1948) [see also the more general Stevens’ law (Stevens, 1957)]. The prediction is that participants will take longer to make decisions and will commit more errors when the JND is small as compared to when it is large. Indeed, discrimination of physical properties such as size, length, weight, as well as more subjective properties such as value, are reported to become slower and more inconsistent when the scale of the discriminated property is increased but the difference is held constant (Busemeyer et al., 2019; Killeen et al., 1993; Marks & Algom, 1998). At the neural level, divisive normalization between neurons has been proposed as a plausible mechanism that can explain such behavioral and neuronal responses to different stimulus strengths (Carandini & Heeger, 2012). However, it has also been shown that certain stimulus properties do not follow the Weber-Fechner law. For example, properties that lack an absolute zero, such as estimates of positions in space, have been reported to show a lack of increase in the deviation as the size of the target object increases (Ganel et al., 2008; Smeets & Brenner, 2008, but see also Utz et al., 2015). Other studies showed that human reaction time decreased as the overall value of choice options increased (Shevlin et al., 2022). This is the opposite of what one would predict if increasing values made their differences less salient, and thus harder to distinguish. In our task, participants made binary choices based on two independent visual attributes, allowing us to investigate whether the response varied as a function of the range of the offered value and attribute types.

## MATERIALS & METHODS

### Participants

Fourteen participants (13 right-handed, 6 females, mean age: 27; range 20-35) participated in the study. Participants 1-3 were colleagues of the authors who volunteered during the pilot study and were not offered monetary compensation. Participants 4-14 signed a consent form prior to the study, and were offered monetary compensation of $20 plus an additional $5-9 based on their final score. All participants had no known neurological conditions and had normal or corrected to normal vision. At the beginning of the session, participants were given verbal instructions describing the task and performed a practice session to familiarize themselves with the task. The practice session lasted until they made 10 consecutive correct choices (i.e., choosing the target with a higher score), and verbally reported that they felt ready. The experimental paradigm was approved by the Comité d’éthique de la recherche en santé of the Université de Montréal.

### Experimental setup

Participants 1-2 performed the task using a standard computer mouse to control a cursor on a screen. The remaining participants performed the task as follows: They were comfortably seated at a planar digitizing tablet (CalComp Drawing Board IV) on which they performed horizontal arm movements by moving a vertically oriented cylindrical handle with their right hand (Figure 1A). The handle contained a wireless digitizing stylus which sent its location to the tablet at 125 Hz. An LCD computer monitor was suspended above the tablet and half-way between them, a semi-silvered mirror that reflected the stimuli displayed on the monitor. The handle with the stylus was placed between the mirror and a digitizing tablet. The monitor, the mirror and the digitizing tablet were equally spaced, creating an illusion that the visual stimuli on the monitor floated at the height of the tablet. During the task, a cross cursor was displayed over the handle to provide continuous feedback about its location.

**Figure 1.**
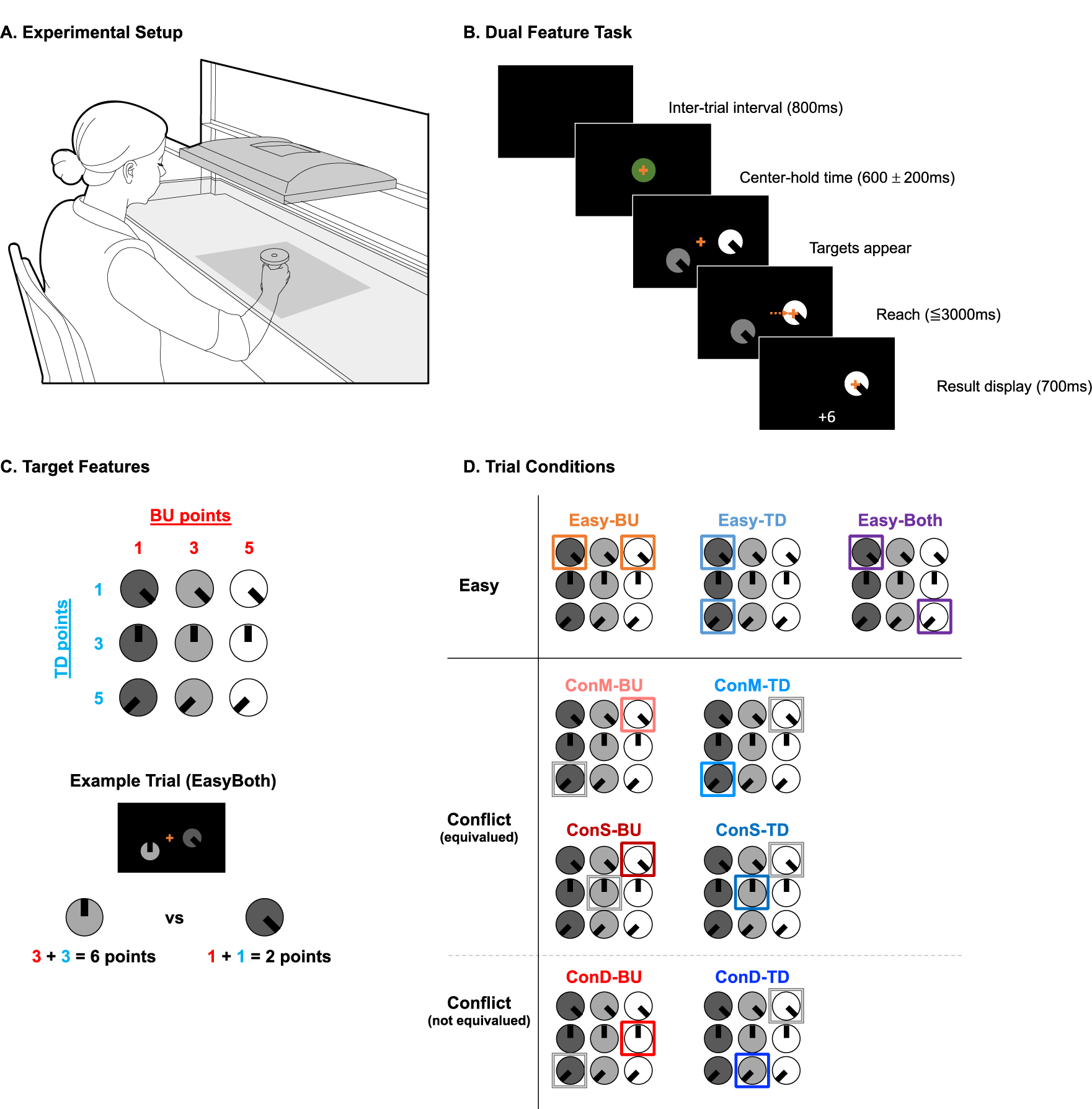
Dual Feature Task. **(A)** Experimental setup, with the participant sitting at the tablet and holding the cursor. **(B)** Progression of the task. After an inter-trial interval, a trial was initiated by moving the cursor (orange cross) into the center target in the middle of the screen. After a variable delay, one or two targets appeared on the right and left bottom of the screen and the center target disappeared. Once the participant chose a target by moving the cursor into it, the unchosen target disappeared and the obtained points were displayed. **(C)** Target Features. The bottom-up (BU) feature was based on three levels of brightness of the target. Targets were given 1, 3 or 5 points if they were dim, medium-bright and bright, respectively. The top-down (TD) feature was based on the orientation of an overlaid line. Targets were given 1, 3 or 5 points if the line was at 4:00, 12:00 and 8:00, respectively. **(D)** Trial conditions and example target combinations. Top, from left to right. Easy-BU (example in orange): targets vary only in BU features. Easy-TD (example in cyan): targets vary only in TD features. Easy-Both (example in violet): targets vary in both features, and both features favor the same target. Bottom, from left to right. Conflict: targets vary in both features, but the BU and TD features favor opposite targets. Two of the conflict conditions, ConM and ConS, consisted of equivalued targets, while ConD consisted of targets with different values. In all conflict conditions, trials were subdivided with a suffix of-BU and -TD based on which feature of the target chosen by the participant had the higher score (these are indicated by a colored frame). ConM (Conflict-Mirror, examples in salmon and pale blue): Conflict trials in which one target’s BU point is the same as the other target’s TD point and vice versa. ConS (Conflict-Same, examples in dark red and dark blue): Conflict trials in which the targets are equivalued, but the BU and TD points did not mirror each other. ConD (Conflict-Different, examples in vivid red and vivid blue): Conflict trials in which the targets had different values. Twin: two identical targets (not shown).

At the beginning of each trial, a circular target (3 cm diameter) appeared in the center of the workspace (Figure 1B). Participants initiated a trial by moving the cursor into this center circle. After a variable delay (600±200 ms, uniform distribution), two circular targets appeared at 0- and 240-degree positions with respect to the center circle, 7.5 cm away, center to center. This choice of targets was used in order to reduce the difference in the biomechanical costs of the associated reaching movements (Cos et al., 2011; Hogan, 1985). Participants then moved the cursor into their target of choice, earning the value of that target toward their total score for the session. The value of each target was indicated by two visual features (Figure 1C). The bottom-up (BU) feature consisted of three possible levels of brightness, such that the brightest target was worth 5 points, a dim target was worth 3 points, and the darkest target was worth 1 point. The top-down (TD) feature consisted of three possible orientations of an overlaid line, such that a line pointing at eight o’clock was worth 5 points, twelve o’clock was worth 3 points, and four o’clock was worth 1 point. The sum of the BU and TD points was earned upon a successful reach to a given target, which was converted into a monetary amount at the end of the session. Once the cursor hit one of the targets, the non-reached target disappeared as feedback confirming the choice. At the end of each trial, the points obtained from the chosen target were added to the total points accumulated, and displayed for 700 ms. The maximum reaction time allowed was 2000 ms, and the maximum movement time was 1000 ms. The inter-trial interval was 800 ms. On average, participants completed 913 trials over the course of 60.4 minutes (range: 684-1085 trials, 41.8-85.5 minutes).

At the end of the session, participants 4-14 received $20 for their participation, plus an additional $5-9 based on the points earned (mean $27.27). The additional amount was determined based on the deviation from the estimated average score, such that the first tercile above the estimated average score would yield an additional $7.00, and the second tercile $8, etc.

### Trial conditions

Each trial was classified as one of 9 conditions based on the features of each target and the participant’s choice (Figure 1D). When the targets varied in brightness (i.e., BU feature) but not in their line orientation, the trial was labelled **Easy-BU**, because the choice was easy (one target is better than the other) and the better target was indicated by the BU feature. When the targets varied in line orientation (i.e., TD feature) but not in their brightness, the trial was labelled **Easy-TD**, as the better target was indicated by the TD feature. When the targets varied in both BU and TD features and both favored the same target, the trial was labelled **Easy-Both**. When the targets were identical, the trial was labelled **Twin**. When the targets varied in both features, but the features favored opposite targets (i.e., one target had better BU feature while the other target had better TD feature), the trial was considered a Conflict trial. These were further categorized as follows: In Conflict-Mirror (**ConM**) trials, the targets’ total scores were equal and the distribution of the feature scores were inverted (e.g., one target had 3 BU points and 1 TD point, while the other had 1 BU point and 3 TD points). In Conflict-Same (**ConS**) trials, the targets’ scores were equal and the distribution of the feature scores were not inverted (e.g., one target had 5 BU points and 1 TD point, while the other had 3 BU points and 3 TD point). In Conflict-Different (**ConD**) trials, the features were in conflict but the target scores were *not* equal. Furthermore, we subcategorized the Conflict trials based on the choice made: if the participant chose the target with the better BU feature, the trial was marked with the suffix BU (i.e., **ConM-BU, ConS-BU, ConD-BU)**, and if the target with the better TD feature was chosen, it was marked with the suffix TD (i.e., **ConM-TD, ConS-TD, ConD-TD)**. As explained below, this was motivated by interest in comparing the timing of choices driven by BU versus TD features.

An average session consisted of 15.7% EasyBU, 15.8% EasyTD, 15.6% EasyBoth, 8% Twin, 14.9% ConM, 10.1% ConS, and 20% ConD trials, all randomly interleaved.

### Behavioral Analysis

Reaction time (RT) was defined as the duration between the stimulus onset and the movement onset. Raw movement velocity was filtered using a 6^th^ order Butterworth low-pass filter with a cut-off frequency at 10 Hz. Movement onset was detected when the filtered velocity exceeded 1.5 cm/second and reached the velocity peak of the trial. Based on the reaching trajectory, each trial was assigned an “Initial Choice” and a “Final Choice”. The Initial Choice was assigned to a target when the cursor moved toward it during the first 100 ms after leaving the center and the linear trajectory estimate fell within 0.6 cm from that target. When a trial did not meet these criteria, the experimenter assigned the Initial Choice via visual inspection. The Final Choice was assigned based on the target which the cursor eventually reached. In a subset of trials, there was a clear change in the reaching trajectory, indicating that the participants changed their mind mid-reach. Such “Change-of-Mind” (CoM) trials were assigned a secondary RT, either at the time when the filtered velocity fell below half of the peak velocity of the initial movement, or when the velocity was at the lowest before it started to increase again as the participant started reaching toward the final target.

Trials were considered outliers when the RT was more than 3 standard deviations away from the median and were excluded from further analysis. When two targets varied in their scores (i.e., Easy and ConD trials), only trials in which participants chose the target with a higher score were included, as the “incorrect” choices may be due to lapses of attention or the participant misinterpreting the features. This resulted in an exclusion of 1449 or 11.5% of trials (mean per participant: 11.9%, range: 5.2-25%). The excluded trials mainly consisted of ConD trials (61%), followed by Easy-TD trials (21.9%), Easy-BU trials (9.2%) and Easy-Both trials (6.4%).

### Statistical Tests

To analyze the difference in the reaction time (RT) distributions across conditions at the individual level, we performed one-way ANOVAs with condition and target positions as factors. Because our task contained free-choice trials (e.g., Twin and Conflict), the number of trials per condition varied across participants. To account for this, we fitted a linear mixed-effects model prior to ANOVA. Target position was included to verify if participants had any directional bias. To analyze the effects of condition at the population level, we converted each participant’s RTs into z-scores by subtracting that participant’s mean RT across all conditions and dividing by the standard deviation across all conditions. This allowed us to combine data from participants with different mean RTs to analyse the order in which different conditions were completed, as we were interested in the relative rather than the absolute values of the RTs across conditions. We then averaged the z-scored RTs across participants to obtain a single mean RT per condition and performed univariate repeated measures ANOVA using a custom MATLAB function (Caplette, 2023). The *p-*values were adjusted with Greenhouse-Geisser to account for the distortion due to the violation of sphericity (Mauchly’s test *χ*^2^(27)=82.60, *p*<0.001). To compare between conditions, we ran post hoc pairwise t-tests. To take multiple comparison into account, the critical value was adjusted using the Bonferroni method. Results were considered significant at *p* < 0.05. All data were processed in MATLAB R2014b.

## RESULTS

### Reaction time distributions varied across conditions

We hypothesized that the BU and TD features would take separate visual pathways to be preferentially processed in different brain regions at different latencies. Therefore, we predicted that different conditions would have different RT distributions within and across participants. Specifically, we predicted that decisions in favor of BU would have a shorter RT distribution from that of TD, especially when the two targets were worth equal values.

First, we analyzed the within-participant effects by focusing on three example participants whose RTs were short (median RT<500 ms), medium (median RT between 500 and 1000 ms), and long (median RT>1000 ms). To analyze the effects of conditions and target positions on the individual participants’ RT distributions, we fitted a linear mixed-effect model and performed a one-way ANOVA with condition and target position as factors. The example participant with short RTs (participant 1, median RT: 316 ms, Figure 2A) had different RTs for different trial conditions (*F*(7,480)=4.12, *p*<0.001). There was also an effect of target position (*F*(1,480)=8.34, *p*=0.004). There was no interaction between the condition and position (*F*(7,480)=0.75, *p*=0.63). The example participant with medium RTs (participant 14, median RT: 656 ms, Figure 2B) had different RTs for different trial conditions (*F*(7,760)=11.74, *p*<0.001). There was no effect of position (*F*(1,760)=0.04, *p*=0.85) nor interaction between condition and position (*F*(7,760)=0.05, *p*=0.05). The effect of condition was observed in all other participants with medium RTs. The example participant with long RTs (participant 5, median RT: 1221 ms, Figure 2C) had different RTs for different conditions (*F*(7,737)=25.52, *p*<0.001). There was no effect of position (*F*(1,737)=3.00, *p*=0.08), but there was an interaction between condition and position (*F*(7,737)=6.61, *p*<0.001). This participant was the only one whose RTs were longer than 1000 ms. Three participants were excluded from within-subject analysis due to the lack of trials in certain conditions (participant 8 made no rightward choices in ConS-TD, participant 9 made no leftward ConS-TD choices, and participant 12 made no ConS-BU choices). In summary, 10 out of 11 participants showed an effect of condition and 3 out of 11 participants showed an effect of position (data not shown).

**Figure 2.**
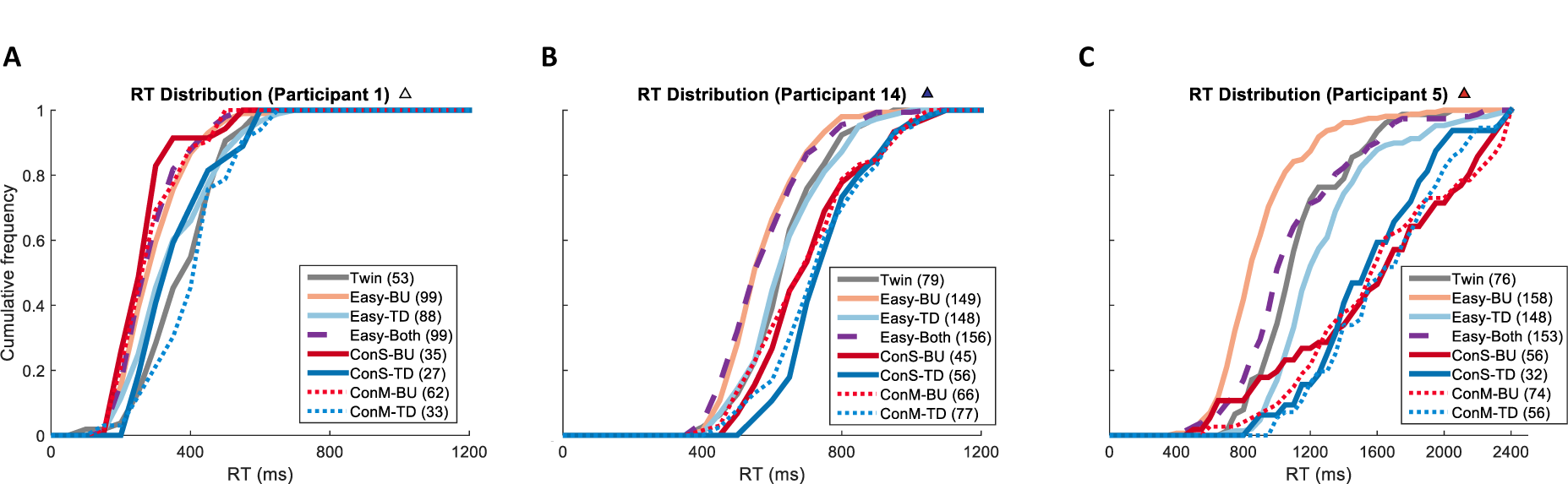
Cumulative distributions of RTs. **(A)** Example participant whose median RTs were short (<500 ms). **(B)** Example participant whose median RTs were medium (between 500 and 1000 ms). **(C)** Example participant whose median RTs were long (>1000 ms). Line colors indicate different conditions. The number of trials of each condition is indicated in parentheses. Each time bin is 50 ms.

To analyze the effects at the population level, we z-scored individual participant’s RTs (see cumulative distributions in Figure 3A) and calculated one mean RT per condition per participant. A univariate repeated measures ANOVA showed an effect of condition (*F*(7,84)=11.32, *p*<0.001). Post hoc pairwise t-tests with multiple comparison correction (see Statistical Tests for details) revealed a significant difference between Easy-BU (z-scored mean *M*=-0.299, *SD*=0.274) and Easy-TD (*M=*0.136, *SD*=0.246) conditions (*p*=0.004); Easy-BU and Twin (*M=*0.251, *SD*=0.285) conditions (*p*<0.001); Easy-BU and ConS-TD (*M=*0.6, *SD*=0.371) conditions (*p*<0.001); Easy-BU and ConM-TD (*M=*0.522, *SD*=0.387) conditions (*p*<0.001); Easy-TD and Easy-Both (*M=*-0.29, *SD*=0.162) conditions (*p*<0.001); Easy-TD and ConS-TD conditions (*p*<0.001); Easy-TD and ConM-TD conditions (*p*=0.007); Easy-Both and Twin conditions (*p*<0.001); Easy-Both and ConS-TD conditions (*p*<0.001); and Easy-Both and ConM-TD conditions (*p*<0.001). In summary, at the population level there were significant RT differences among Easy trials and between Easy and Conflict-TD trials, but no significant differences within Conflict trials were observed.

**Figure 3.**
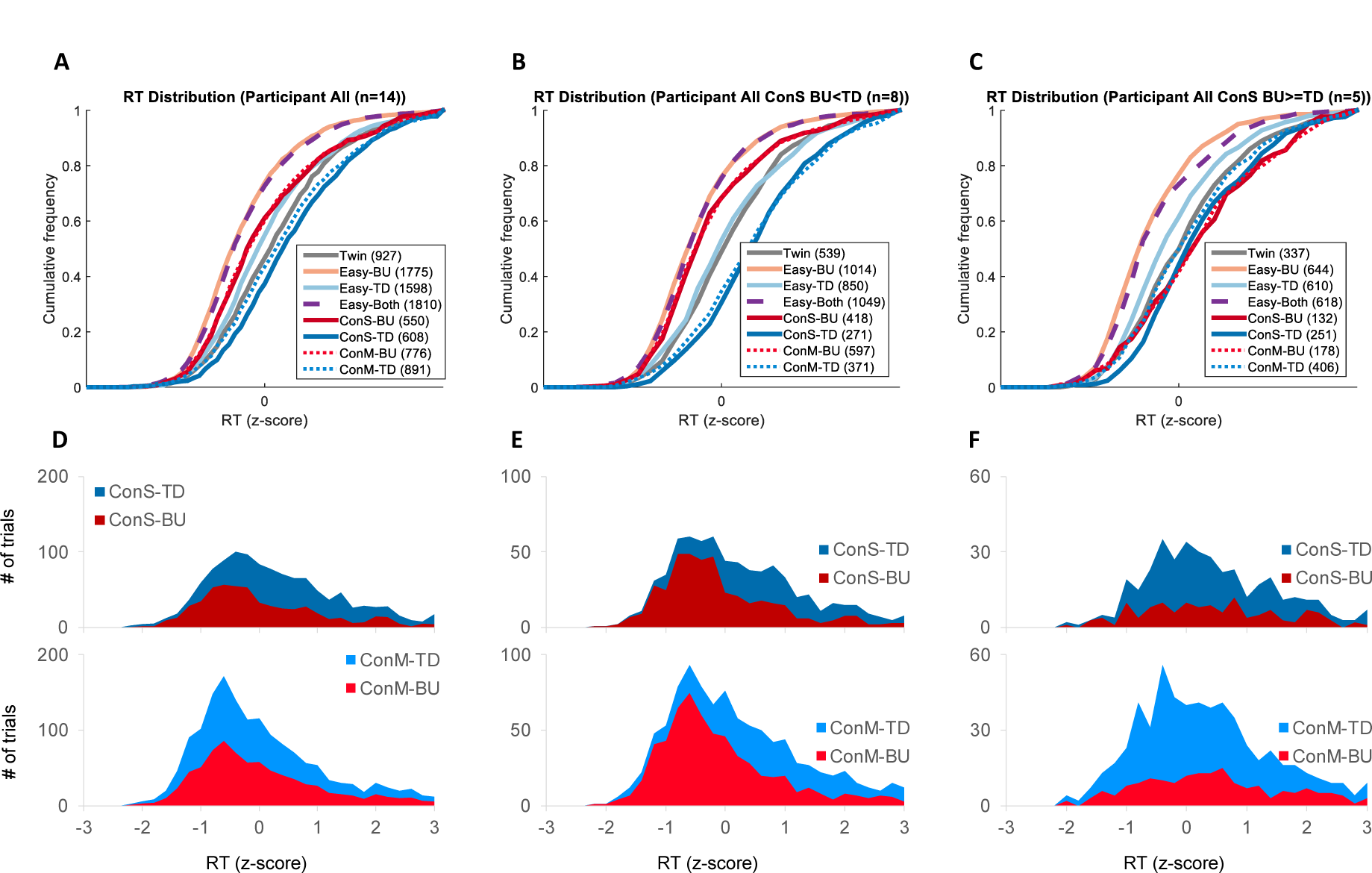
Cumulative distributions of z-scored RTs of all participants. **(A)** Z-scored average RTs of all participants. **(B)** Z-scored average RTs of participants whose RT for ConS-BU was shorter than that of ConS-TD (N=9). **(C)** Z-scored RTs of participants whose RT for ConS-BU was equal or longer than that of ConS-TD (N=4). In all groups, Easy-BU (orange) and Easy-Both (dashed purple) had the shortest RT. **(D-F)** Number of BU and TD choices during ConS (top) and ConM (bottom) trials, plotted as a function of z-scored RT. Subgroups are the same as in plots A-C.

Not all participants exhibited the same trends as the average shown in Figure 3A. In particular, some participants’ RTs were much shorter for ConS-BU compared to ConS-TD trials. We thus subdivided the participants into two groups based on the median RTs and Wilcoxon rank sum tests. Eight participants displayed significantly shorter RTs for ConS-BU than ConS-TD, while five participants did not meet statistical significance (data not shown). For the first group (Figure 3B), a univariate repeated measures ANOVA showed an effect of condition (*F*(7,49)=17.71, *p*<0.001). Post hoc pairwise t-tests with multiple comparison correction revealed a significant difference between Easy-BU (*M=*-0.316, *SD*=0.233) and Twin (*M=*0.244, *SD*=0.257) conditions (*p*=0.009); Easy-BU and ConS-TD (*M=*0.756, *SD*=0.401) conditions (*p*=0.019); Easy-BU and ConM-TD (*M=*0.698, *SD*=0.355) conditions (*p*=0.013); Easy-TD (*M=*0.224, *SD*=0.291) and Easy-Both (*M=*-0.382, *SD*=0.169) conditions (*p*=0.017); Easy-TD and ConS-TD conditions (*p*=0.046); Easy-Both and Twin conditions (*p*=0.002); Easy-Both and ConS-TD conditions (*p*=0.009); Easy-Both and ConM-TD conditions (*p=0.01*); ConS-BU (*M=*-0.119, *SD*=0.241) and ConS-TD conditions (*p*=0.012); and ConS-BU and ConM-TD conditions (*p*=0.029). In summary, this subgroup exhibited significant differences similar to the trends observed in the overall population. For the group that did not show RT differences between ConS-BU and ConS-TD trials (Figure 3C), a univariate repeated measures ANOVA showed an effect of condition (*F*(7,28)=3.69, *p*<0.006). Post hoc pairwise t-tests with multiple comparison correction revealed significant difference between Easy-Both (*M=*-0.291, *SD*=0.172) and ConS-TD (*M=*0.45, *SD*=0.16) conditions (*p*=0.037). This subgroup did not exhibit the same trend as the overall population. In conclusion, when participants had shorter RTs for ConS-BU than ConS-TD, their RT distributions varied across different conditions, suggesting that they were processing BU and TD features at different latency. In contrast, participants with similar RTs for ConS-BU and ConS-TD conditions completed most of the conditions at similar latency, suggesting that they may have been integrating all information prior to making a choice.

Table 1 shows the medians of the z-scored RTs across participants, as well as that of the two subgroups. Note that there is an inherent skew in reaction time distributions, especially for slower participants, where the right tail of the RT distributions tends to be longer than the left. Thus, when data are averaged together, the RT values from slow participants can dominate those from fast ones. Since here we are primarily interested in comparing differences across trial types, we focus most of our conclusions on the results of z-scored data. At a glance, the trends between the whole population and the second subgroup are similar. In the second subgroup, RT for ConS-BU was shorter than that of ConS-TD, and their z-scored RTs for Easy-BU, Easy-Both, ConS-BU and ConM-BU were below zero, suggesting that BU-driven choices were made faster than the global mean. In contrast, the third subgroup, whose RT for ConS-BU was not significantly different than that of ConS-TD, all conflict conditions had positive RTs, while only Easy-BU and Easy-Both were below zero.

**Table 1.** Z-scored mean RT of **(A)** all participants, **(B)** participants whose RT for ConS-BU was shorter than that of ConS-TD, and **(C)** participants whose RT for ConS-BU was not significantly different from that of ConS-TD.

|  | All participants<br>z-scored mean RT (SD) |  | ConSBU < ConSTD<br>z-scored mean RT (SD) |  | ConSBU ≥ ConSTD<br>z-scored mean RT (SD) |  |
| --- | --- | --- | --- | --- | --- | --- |
| Easy-BU | -0.30 | (0.27) | -0.32 | (0.23) | -0.34 | (0.35) |
| Easy-TD | 0.14 | (0.25) | 0.22 | (0.29) | 0.02 | (0.11) |
| Easy-Both | -0.29 | (0.16) | -0.31 | (0.17) | -0.29 | (0.17) |
| Twin | 0.25 | (0.29) | 0.24 | (0.26) | 0.19 | (0.34) |
| ConS-BU | 0.14 | (0.48) | -0.12 | (0.24) | 0.56 | (0.48) |
| ConS-TD | 0.60 | (0.37) | 0.76 | (0.40) | 0.45 | (0.16) |
| ConM-BU | 0.15 | (0.56) | -0.07 | (0.32) | 0.48 | (0.77) |
| ConM-TD | 0.52 | (0.39) | 0.70 | (0.35) | 0.35 | (0.30) |

### Reaction time was generally longer during Conflict trials, and when choosing a TD-favored target

To visualize the global trend, we generated a scatterplot of each participant’s mean RT in different trial conditions (Figure 4). Within Easy trials, 10/14 participants were faster in Easy-BU than Easy-TD (two sample t-test at *p*<0.05, Figure 4A). During ConS trials, 6/13 participants were significantly faster when choosing BU-favored targets while 1/13 was faster when choosing TD-favored targets (Figure 4B). This comparison could not be performed for participant #12 because that person never chose the BU target in ConS trials. During ConM trials, 5/14 participants were faster when choosing BU-favored targets while 1/14 was faster when choosing TD-favored targets (Figure 4C). When all Easy trials and all equivalued Conflict trials are collapsed together, 8/14 participants were faster in Easy trials (Figure 4D). Similarly, 12/14 participants were faster in Easy trials than Twin trials (Figure 4G). Between Twin and all equivalued Conflict trials, 3/14 participants were faster in Twin whereas 6/14 were faster in Conflict trials (Figure 4H). During ConD trials, 8/14 were faster when choosing BU-favored targets, while 1/14 was faster when choosing TD-favored targets (Figure 4I). There was no systematic clustering observed in the RT distributions.

**Figure 4.**
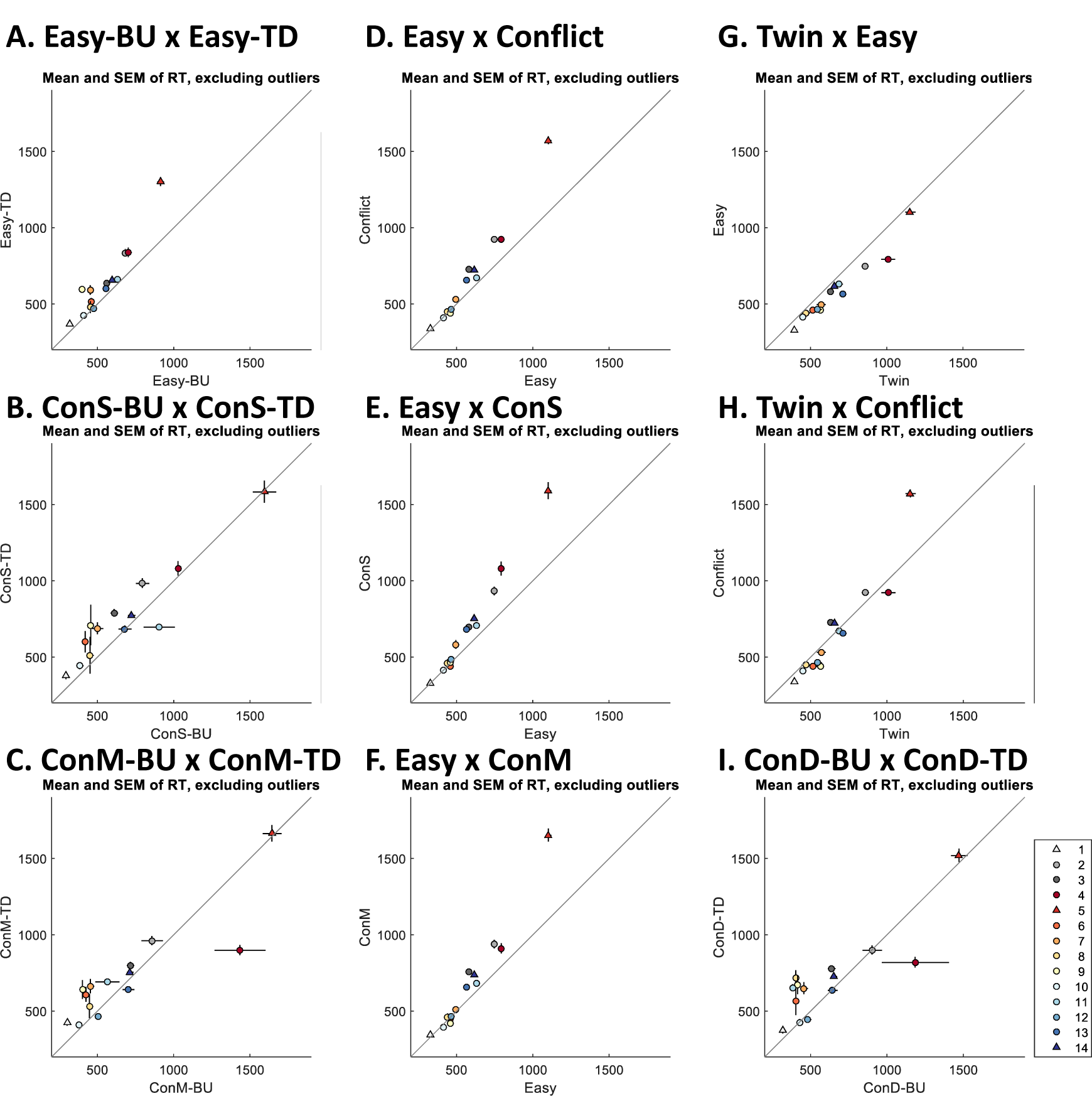
Scatter plot of mean RT across trial conditions. Circles and triangles indicate the mean reaction time of each participant (identified as shown in the legend at bottom right). The standard error of the mean is indicated by the overlaid lines along the respective dimensions. **(A-C)** Comparison of RT between BU and TD features for Easy, ConS and ConM trials. **(D-F)** Comparison of Easy and Conflict trials. **(G-H)** Comparison of Twin trials against Easy and Conflict trials. **(I)** Comparison of ConD-BU and ConD-TD trials. The detailed data of the participants indicated by triangles, whose median RTs were short (<500 ms, light gray), medium (between 500 and 1000 ms, dark blue) and long (>1000 ms, red), are further discussed above.

### Participants made faster choices when choosing between two high-valued targets

The Dual Feature task was designed as a value-based decision task. However, due to the nature of the stimuli we used, one could say that it is also a perceptual discrimination task. Importantly, while the BU feature has a linear component (dark to bright), the TD feature was more arbitrary, and the discrimination between different line orientations was not comparable to the discrimination between different brightness levels. This allowed us to test whether the Weber-Fechner law, which postulates that perceptual discrimination becomes more difficult (and therefore slower) as the amplitude of the discriminated stimuli increases, applies when the discriminated stimuli are associated with reward magnitude. Specifically, we analyzed the reaction time of choices between two high-valued targets versus two low-valued targets by comparing the reaction times when the choice was based on the attribute whose stimulus strength was linearly correlated with the reward magnitude (i.e., Easy-BU) or not (i.e., Easy-TD).

We compared the RT and choice accuracy of Easy trials that offered two of the highest and two of the lowest possible values (Figure 5). In high reward trials, the choice is between two high-value targets, one worth 10 points (5 BU + 5 TD points) and another worth 8 points (Easy-BU: 5 BU and 3 TD, or Easy-TD: 3 BU and 5 TD). In low reward trials, the choice is between a target worth 4 points (Easy-BU: 3 BU and 1 TD, or Easy-TD: 1 BU and 3 TD) and another worth 2 points (1 BU and 1 TD). In Easy-BU trials, there was no significant difference in RT between low (*M=*594.6, *SD*=251.3) and high (*M=*566.2, *SD*=252.1) reward offers (*p*=0.15, two-sample Kolmogorov-Smirnov test). The accuracy was better for low (*M*=0.95, *SD*=0.09) than high (*M*=0.86, *SD*=0.11) trials (*t*(26)=-2.27, *p*=0.031, two-sample t-test) offers. By contrast, in Easy-TD trials, RT was shorter when the offered rewards were higher (*M=*777.94, *SD*=334.95) than lower (*M=*604.29, *SD*=290.09, *p*<0.001). The accuracy was not significantly different between low (*M*=0.78, *SD*=0.22) and high (*M*=0.87, *SD*=0.12) offers (*t*(26)=1.38, *p*=0.17).

**Figure 5.**
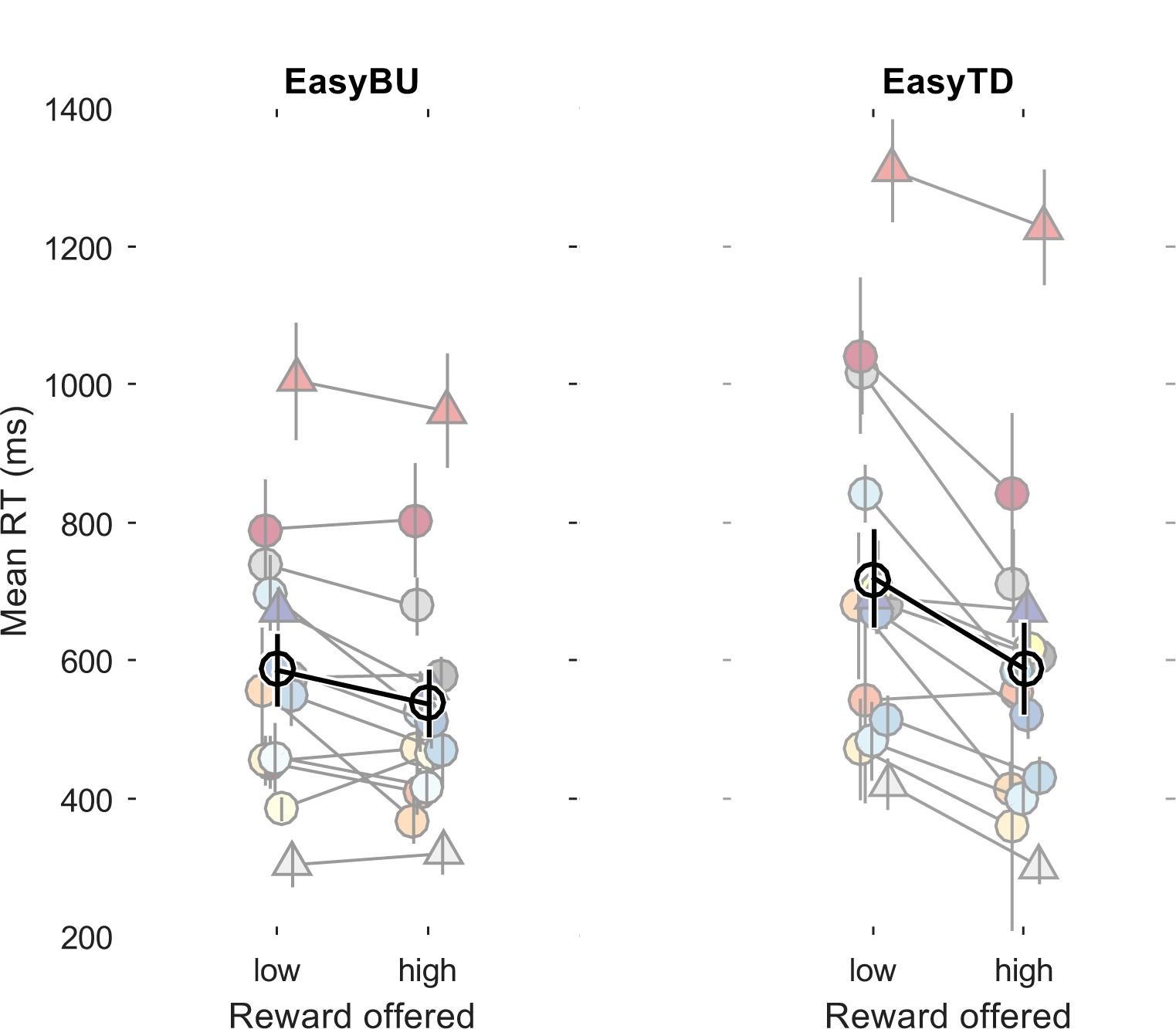
Mean RTs grouped by offered reward magnitude. RT did not increase when the offered reward was larger. Different colours correspond to different participants. The triangles represent the example participants from Figure 2, whose mean RTs were short (<500 ms, light gray), medium (between 500 and 1000 ms, dark blue) and long (>1000 ms, red). Black open circle with black overlaid lines indicates the mean RT of all participants with standard errors of the mean.

The mean RT of Easy-BU trials did not vary between high and low offers, though participants made more mistakes when the offers were higher. In fact, the accuracy for low offer Easy-BU trials were almost at the ceiling. The mean RT for Easy-TD trials was shorter for higher offers and the accuracy was comparable.

In Easy-BU trials, the observed decrease in the accuracy for high offers may be due to the Weber-Fechner law in brightness discrimination, although there was no difference in the RTs between high and low offers. In Easy-TD trials, the presence of the highest BU features reduced the latency of the TD-based choices, although the accuracy was unaffected. Overall, our results are not compatible with Weber-Fechner law, which predicts both reduction in the accuracy and increased latency for higher offers. The reduction in the latency in high offer Easy-TD trials is similar to what has been reported in previous studies (Shevlin et al., 2022).

### Change of mind happened mostly as a switch from the BU-favored to the TD-favored target

All participants showed some changes-of-mind (CoM), particularly during conflict trials, indicated by an abrupt change in the movement trajectory (see examples in Figure 6A). Participant 12 had no ConS-BU trials because they changed their mind in all of those trials, making them ultimately end up as ConS-TD trials. Out of 3467 equivalued conflict trials, 174 trials (5%) contained a change-of-mind (68 in ConS trials, 106 in ConM trials). In ConS, 57/68 (84%) changes were from the BU-favored to the TD-favored target (Figure 6B, left). In ConM, 83/106 (78%) were from the BU-to the TD-favored target (Figure 6B, right). Thus, most changes of mind happened as a switch from BU to TD.

**Figure 6.**
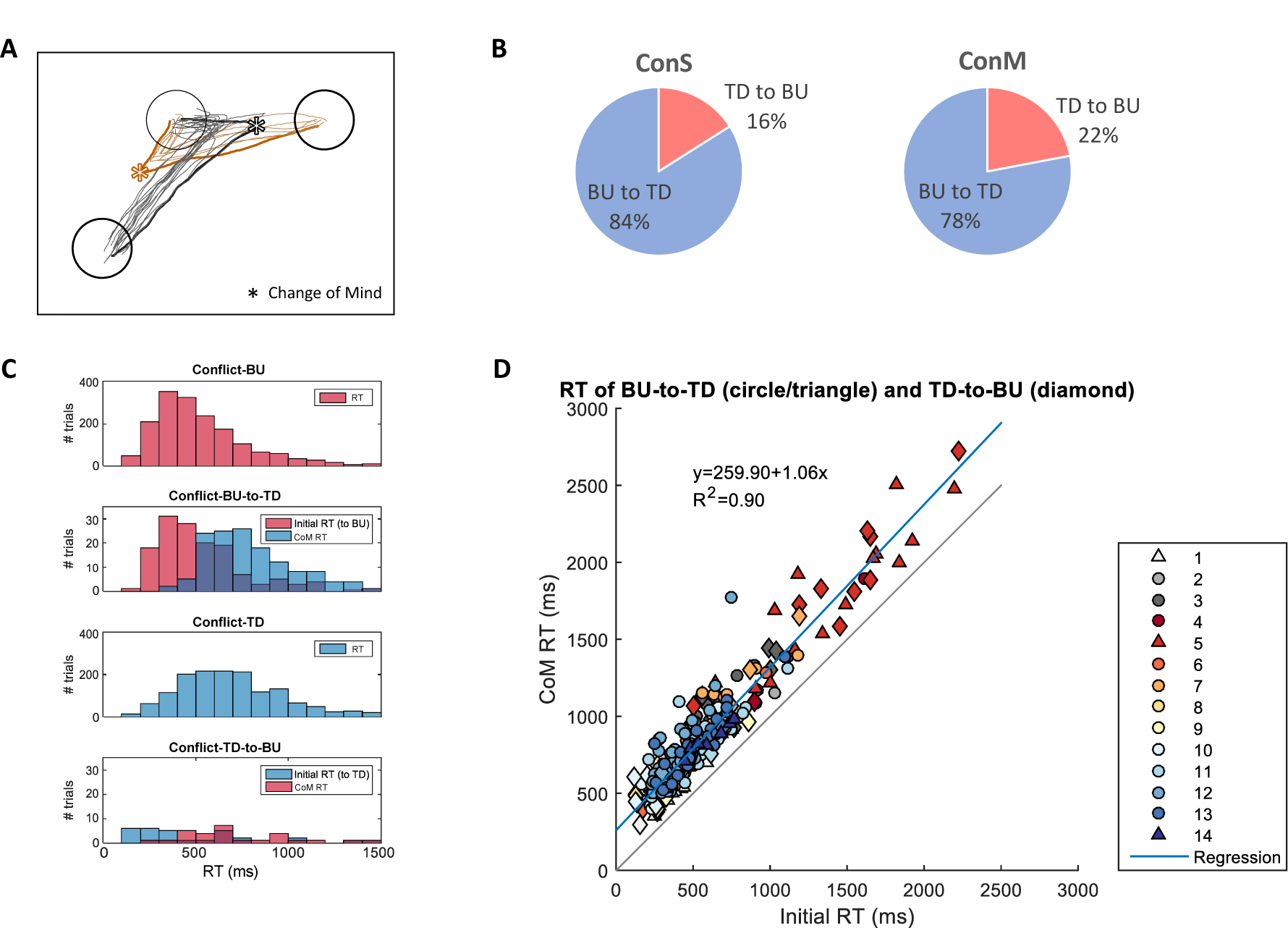
Change of Mind (CoM) trials. **(A)** Sample trajectories of change-of-mind trials. Black and brown lines are trajectories to bottom left and right targets, respectively. Stars indicate the point in which CoM was assigned for two example trials (thick lines). **(B)** In ConS trials, 84% of CoM were from BU to TD. In ConM trials, 78% of CoM were from BU to TD. **(C)** Reaction time distributions of conflict trials with and without CoM. TOP to BOTTOM: Conflict-BU trials without CoM (i.e., straight trajectory). Conflict trials in which the initial trajectory was toward the BU-favored target (red RT), but finally reached the TD-favored target (blue CoM RT). Conflict-TD trials without CoM. Conflict trials in which the initial trajectory was toward the TD-favored target (blue RT), but finally reached the BU-favored target (red CoM RT). **(D)** Scatter plot of initial and CoM RT of individual trials. Different colors correspond to different participants. Circles and triangles: CoM trials changing from BU to TD. Diamonds: initially CoM trials changing from TD to BU. Blue line: Linear regression, p<0.001, R^2^=0.91.

We tested whether the CoM was a result of participants correcting their premature decisions by comparing the RTs of Conflict trials with and without CoM (Figure 6C). The initial RT of BU-to-TD CoM trials (*M*=563.66, *SD*=327.28) and the RT of non-CoM Conflict-BU trials (*M*=588.92, *SD*=392.73) were not significantly different (*t*(1880)=0.75, *p=*0.4, two-sample t-test, compare red distributions in the two top rows of Figure 6C). The initial RT of TD-to-BU CoM trials (*M*=608, *SD*=521.13) was faster than the RT of non-CoM Conflict-TD trials (*M*=744.15, *SD*=364.8, *t*(1576)=2.1, *p*=0.036, compare blue distributions in two bottom rows of Figure 6C). There was no statistical difference between the initial RTs of the BU-to-TD and TD-to-BU CoM trials (*t*(176)=-0.62, *p=*0.54). As shown in Figure 6D, the timing of the CoM was not constant with respect to target onset, but instead followed the initial RT by approximately 266 ms (regression slope = 1.04, R^2^=0.91, *p<0.001*).

## DISCUSSION

We investigated whether humans make value-based decisions by fully integrating all available information into a unified measure of subjective value, or whether such decisions involve a more distributed process whereby separate sources of information can themselves compete. Participants chose one of two reachable targets whose value was based on two visual attributes, the brightness of the target (a bottom-up cue, BU), and an oriented line overlaid on the target circle (a top-down cue, TD). We predicted that, if the BU and TD attributes in our task are always converted into representation of reward value and decisions are made by comparing the total value of each target, RT distributions would be faster when only one feature is different (Easy-BU and Easy-TD trials) and slower in more complex conditions where both features differ (Easy-Both and Conflict trials), but there would be no difference between Conflict-BU versus Conflict-TD trials. Under this “integrated competition” hypothesis, the differences in RTs between different trial conditions would be due to the time it takes to process the BU and TD features. If we assume that the comparison process is bypassed when a feature is identical across the targets, then we would predict that RTs in Easy-BU trials would be the fastest, followed by Easy-TD, then followed by Easy-Both and Conflict conditions. In this view, Conflict trials might be expected to be slower than Easy-Both trials, as a decision may require recruiting an additional process that resolves the conflict. Most importantly, however, identifying Conflict trials could only be done after both features were processed, so there should be no difference between the RT distribution of Conflict-BU versus Conflict-TD trials.

Alternatively, if the BU and TD attributes in our task are processed in different neural substrates, and the decision could be made prior to a complete integration at the level of total reward value, then differences in the RTs should reflect which attribute was used to decide, especially during trials with equally valued targets. That is, a “distributed competition” hypothesis predicts that decisions based on the BU attribute would be faster than those based on TD attributes in both Easy and Conflict trials, and choices in favor of the BU attribute (Conflict-BU) would be more frequent in early decisions whereas those in favor of the TD attribute (Conflict-TD) would be more frequent in later decisions.

We found that RTs varied across trial types and that the order of the conditions was not strictly congruent with the complexity of the features. Notably, all participants were fastest in Easy-BU and Easy-Both trials, which was followed by either ConS-BU and ConM-BU trials or Easy-TD trials, creating two subgroups of participants. If all information was integrated before making the decision, one would expect that the latency for Easy-Both to have been similar to that of Easy-TD, as solving Easy-Both trials should take at least as long as processing the TD feature in Easy-TD trials. Furthermore, in the subgroup of participants with ConS-BU and ConM-BU as the second fastest condition, Easy-TD was the third condition. This suggests that during these types of Conflict trials, participants often chose the BU-favored target before taking the TD feature into account. In this subgroup, the slowest condition was ConS-TD and ConM-TD, suggesting that choosing in favor of the TD feature in Conflict trials takes longer than choosing a left or right target when the two targets are identical (Twin trials). For the other subgroup of participants, Easy-TD and Twin were the second fastest conditions, and ConS-BU, ConM-BU, ConS-TD and ConM-TD conditions were all similar and the slowest. In other words, these participants were faster when solving Easy trials and slower when the choice was ambiguous (i.e., the targets were equal-valued), especially when the features were in conflict. However, the overlap between the RT distributions of Easy-Both and Easy-BU trials and the fact these subjects took longer to solve Easy-TD than Easy-Both are incompatible with decision timing being simply determined by trial complexity. Instead, these results are compatible with the proposal that BU and TD features are processed by different neural mechanisms and the decision can sometimes be made before both processes are complete.

Furthermore, we saw that changes-of-mind in Conflict trials were mostly from the BU-favored target to the TD-favored target. The timing of the change of mind was not constant with respect to target onset, but instead followed the initial RT by approximately 266 ms. Importantly, changes from the TD-favored to a BU-favored target occurred most often when the initial RT was very short, again suggesting that subjects sometimes make choices before integrating all sources of information about value.

Finally, we investigated whether the BU and TD features in our task follow predictions of the Weber-Fechner Law, a psychophysical theory that posits an increase in the just-noticeable-difference proportional to the magnitude of the stimuli. We subdivided Easy-BU and Easy-TD trials based on the magnitude of offered reward, and compared the RTs to see if deciding between two large offers took longer than deciding between two small offers, as would be expected if the just-noticeable-difference was smaller and thus took longer to discriminate. We found that the magnitude of the reward offered did not influence RT in Easy-BU trials, but higher offers negatively affected choice accuracy. During Easy-TD trials, participants were faster when choosing between two high reward options than when they were choosing between two low reward options, whereas the choice accuracy was similar between the two. This was partially in line with previous results that argued against diminishing value sensitivity in value-based decisions (Shevlin et al., 2022).

Taken together, our results suggest that despite indicating the same type of information (i.e., the reward value of a target), the two visual features we chose were not simply integrated into a “common currency”. Instead, our data is consistent with the premise that BU and TD features are treated differently, possibly by different neural substrates. The finding that a subset of participants had shorter RT in ConS-BU and ConM-BU trials than in Easy-TD trials suggests that the decision processes that favor BU features can sometimes bypass the processing of the TD feature. That is, a decision-making area in the brain that has received information about the BU feature has the capability to trigger the choice without waiting for the complete integration of information about the TD feature. In summary, our results suggest that action decisions can be made before all information is integrated, supporting the proposal that a hierarchy of within- and between-attribute competitions influences the higher-order choice-level competition (Hunt et al., 2014).

There are many possibilities of where in the brain such competitive processes might unfold. For example, based on functional magnetic resonance imaging (fMRI) data in humans performing multi-attribute multi-alternative choice task, Hunt and colleagues suggested that cells in the intraparietal sulcus signal the between-attribute competition, whereas medial prefrontal cortex signals the between-option competition at the ‘integrated’ value level (Hunt et al., 2014). Other human fMRI studies suggested that the orbitofrontal and ventromedial prefrontal cortex reflect subjective value comparison across categories such as food, money, consumable products and pain (FitzGerald et al., 2009; Kable & Glimcher, 2007; Smith et al., 2010; Talmi et al., 2009). When tasks require incorporation of negative values such as effort, monetary loss, pain or satiation-induced devaluation, ventromedial prefrontal cortex, anterior cingulate cortex, supplementary motor area, amygdala and nucleus accumbens have been suggested to provide the devaluation signal (Basten et al., 2010; Croxson et al., 2009; Gottfried et al., 2003; Klein-Flügge et al., 2016; Talmi et al., 2009). In a seminal study, Buschman & Miller (2007) showed that when monkeys perform a visual pop-out task in which the correct target can be easily discriminated from among distractors in a bottom-up fashion, choice-related activity appears in posterior parietal cortex before frontal regions, but when the task requires a slower serial visual search the opposite pattern of latencies is observed. This suggests that bottom-up decision tasks may be solved by the fast dorsal visual stream (Ledberg et al., 2007; Schmolesky et al., 1998), while more complex tasks require slower processes that combine distinct visual features, implicating the ventral visual stream and its projections to prefrontal regions (Donner et al., 2002; Yan et al., 2016) In our experiment, both of these kinds of processes may be taking place within the context of a single task, even during individual trials. This could make it possible to identify where in the brain the two kinds of relevant cues – bottom-up and top-down – are processed, and whether they compete in a distributed manner or only after being integrated into a unified estimate of subjective value within a single “central executive”. Such studies are underway (Nakahashi et al., 2018; Nakahashi & Cisek, 2016, 2020), but their results are beyond the scope of the current paper.

## ACKNOWLEDGEMENTS

The authors thank Aarlenne Khan for helpful comments on an earlier draft of this manuscript. This work was supported by grants from the Canadian Institutes of Health Research (PJT-166014), the Natural Sciences and Engineering Research Council of Canada (RGPIN/05245), and the Fonds de la recherche en santé du Québec to PC, and scholarships from the Groupe de Recherche sur le Système Nerveux Central and Fonds de Recherche Québec Nature et Technologies to AN.

